# Optimizing the detection of biological signals through a semi-automated feature selection tool

**DOI:** 10.1101/2024.08.07.607073

**Authors:** Gabriel Santos Arini, Luiz Gabriel Mencucin, Rafael de Felício, Luís Guilherme Pereira Feitosa, Paula Rezende-Teixeira, Henrique Tsuji, Alan Pilon, Danielle Rocha Pinho, Letícia Veras Costa-Lotufo, Norberto Peporine Lopes, Daniela Barretto Barbosa Trivella, Ricardo Roberto da Silva

**Affiliations:** Computational Chemical Biology Laboratory, Department of BioMolecular Sciences, School of Pharmaceutical Sciences of Ribeirão Preto, University of São Paulo, Ribeirão Preto 14040-900, Brazil; NPPNS, Department of BioMolecular Sciences, School of Pharmaceutical Sciences of Ribeirão Preto, University of São Paulo, Ribeirão Preto, 14040-900, Brazil; Cellular and Molecular Biology Program, Department of Cellular and Molecular Biology of Ribeirão Preto, School of Medicine, University of São Paulo, Ribeirão Preto, 14049-900, Brazil; Brazilian Biosciences National Laboratory (LNBio), Brazilian Center for Research in Energy and Materials (CNPEM), Campinas, 13083-100, Brazil; Marine Pharmacology Laboratory, Department of Pharmacology, Institute of Biomedical Sciences, University of São Paulo, São Paulo, 05508-000, Brazil; Department of Biochemistry and Organic Chemistry, Institute of Chemistry, Paulista State University, São Paulo, 14800-060, Brazil

**Keywords:** mass spectrometry, data dependent acquisition, chemometrics, untargeted metabolomics

## Abstract

Untargeted metabolomics is often used in studies that aim to trace the metabolic profile in a broad context, with the data-dependent acquisition (DDA) mode being the most commonly used method. However, this approach has the limitation that not all detected ions are fragmented in the data acquisition process, in addition to the lack of specificity regarding the process of fragmentation of biological signals. The present work aims to extend the detection of biological signals and contribute to overcoming the fragmentation limits of the DDA mode with a dynamic procedure that combines experimental and in silico approaches. Metabolomic analysis was performed on three different species of actinomycetes using liquid chromatography coupled to mass spectrometry. The data obtained were preprocessed by the MZmine software and processed by the custom package, RegFilter. RegFilter allowed the coverage of the entire chromatographic run and the selection of precursor ions for fragmentation that were previously missed in DDA mode. Most of the ions selected by the tool could be annotated through three levels of annotation, presenting biological relevant candidates. In addition, the tool offers the possibility of creating local spectral libraries curated according to the user’s interests. Thus, the adoption of a dynamic analysis flow using RegFilter allowed for detection optimization of biological signals, previously absent in the DDA mode. In addition, this workflow enables the creation and search of in-house tailored custom libraries.

## 1 Introduction

Metabolomics offers the possibility of molecular phenotyping and has been applied to diverse fields of knowledge, from biotechnology to environmental chemistry (Kuehnbaum and Britz-McKibbin, 2013). Unlike nucleic acids and even proteins, the metabolites produced in a given biological context help the understanding of rapid spatio-temporal changes triggered by the environment. To harness the metabolic complexity, metabolomics uses several analytical techniques, focusing on mass spectrometry (MS) (Wehrens and Saleck, 2019). Targeted metabolomics focuses on specific metabolites, and its main advantage, besides specificity, is its reproducible quantitative aspect. In contrast, untargeted metabolomics is characterized by the enormous size and complexity of the data sets generated. One of the major advantages of this approach is the possibility of profiling a large number of new molecules, which in turn would have the capacity to reveal unexplored metabolic pathways (Prosser et al., 2014; Banh et al., 2021), establish evolutionary parallels (Nunes et al., 2017; Roach et al., 2021), offer possible insights about what is happening in a given cellular scenario (Zaramela et al., 2021), and present new natural products of biotechnological and pharmacological interest in a wide and diverse range of organisms (Wilke et al., 2021; Kato et al., 2024). Currently, more than 60% of pharmaceutical assets are related directly or indirectly to natural products, i.e. chemical substances produced by living organisms (Newman and Cragg, 2020; Kim et al., 2021).

Often, the data generated in untargeted metabolomics experiments are not explored in their entirety. The recent advances in data sharing, and availability of public spectral data libraries, in platforms such as Metlin (Xue et al., 2020), Metabolights (Yurekten et al., 2024), MoNA (https://mona.fiehnlab.ucdavis.edu/), Metabolomics Workbench (Sud et al., 2016) and GNPS (Wang et al., 2016; Wang et al., 2020) as well as analytical and integrative tools such as Mzmine (Schmid et al., 2023), NAP (da Silva et al., 2018), MolNetEnhancer (Ernst et al., 2019), ChemWalker (Borelli et al., 2023), ModiFinder (Shahneh et al., 2024), and NP^3^ MS Workflow (Bazzano et al., 2024) greatly expanded the potential of metabolomics application. However, the highly redundant and noisy data of mass spectra from complex mixtures is a great challenge. A promising strategy would be to target selected signals, deepening studies within a given context of interest, rationally moving from an untargeted to a targeted approach. This strategy is justified by the fact that many of the detected ions are not fragmented in the mostly used fragmentation mode for untargeted metabolomics: the data-dependent acquisition (DDA) mode (Broeckling et al., 2018; Davies et al., 2021).

Reasons for the preference for DDA in untargeted metabolomics experiments include the availability of optimized tools and workflows for acquiring and processing these data (Stincone et al., 2023; Guo and Huan, 2020), as well as the ready availability of the acquired spectra and their quality (McBride et al., 2023). However, one of the main limitations of using this data acquisition mode in MS^2^ is the non-fragmentation of a number of features (Broeckling et al., 2018). Despite the progress that has been made in terms of increased fragmentation coverage, it is reasonable to assume that much relevant biological information is lost in this process.Moving from an untargeted approach to rationally selecting relevant features in a given context is a logical next step in metabolomics research. The construction of feature selection lists based on objective statistical criteria has been presented as a promising resource in this context (Zhang et al., 2023). However, the proper automation of the process, as well as the ability to store the fragmentation spectra resulting from this selection process, is still a task to be overcome.

In this scenario, the present work aims to introduce an hybrid and dynamic approach which combines experimental and computational procedures, providing a feature selection tool that optimizes the detection of biological signals, and contributes to overcoming the fragmentation limits of the DDA mode and the ability to create in-house spectral libraries.

## 2 Materials and Methods

### 2.1 Collection, extract preparation, and metabolomics analysis

The actinomycetes BRA006, BRA010, and BRA177, from MicroMarin collection (https://www.labbmar.ufc.br/micromarinbr) were cultured in A1 medium [Starch (10 g/L); Yeast extract (4 g/L); Peptone (2 g/L); Sea Water 75%] in a volume of 100 mL in 250 mL Erlenmeyer flasks. The liquid cultures were extracted with ethyl acetate, and the organic phase was dried under pressure and kept at 4°C. For the LC-MS/MS analyses, organic extracts were diluted in methanol and injected at 1.0 mg/mL, 0.5 mg/mL, 0.25 mg/mL, 0.125 mg/mL, and 0.0625 mg/mL concentrations in 1:1 H_2_O:metanol (Bauermeister et al., 2016). The LC-MS/MS analysis itself was conducted in the Acquity UPLC H-Class (Waters, Milford, MA – US) hyphenated with Impact II mass spectrometer (Bruker Daltonics, Billerica – US). The mobile phase (flow 0.3 mL.min-1) consisted of water (A) and methanol (B) acidified with formic acid at 0.1% in the following gradient: 0.0 – 15.0 min (5 – 20% B, curve 6); 15.0 – 30.0 min (20-95% B, curve 6); 30.0 – 33.0 min – (100 B, curve 1); 33.0 – 40.0 min (5% B, curve 1). C18 – Luna (Phenomonex® – 100 mm x 2.1 mm x 2.6 μm) and the temperature adjusted to 35°C. The parameters adjusted for the spectrometer were: end plate offset of 500V; capillary voltage of 4.5kV; nitrogen (N_2_) was used as gas; drying gas flow at 5.0L.min-1; drying gas temperature at 180°C; 4 bar nebulizer gas pressure; positive ESI mode. Spectra (*m/z* 30–2000) were recorded at a rate of 8 Hz. Accurate masses were obtained using a solution of sodium formic acid [HCOO-Na+] at 10 mM as an internal standard.

### 2.2 Computational methods

#### 2.2.1 Raw data conversion and Preprocessing through MZmine

Raw MS files were converted to .mzXML format using MSConvert software (Chambers et al., 2012). Such converted files were pre-processed by MZmine software version 2.53. Data preprocessing in full scan and MS/MS modes was performed separately. The following parameters were used for the full scan mode: (i) Mass detection (MS level 1): mass detector – Centroid; noise level – 1E3; (ii) ADAP: min group size – 4; Group intensity threshould – 1E3; Min highest intensity – 3E3; *m/z* tolerance – 0.001*m/z* or 15ppm; (iii) Chromatogram deconvolution: algorithm – Baseline cut-off; Min peak height – 3E3; Peak duration range: 0 – 15; Baseline level – 1E3; *m/z* center calculation: Median; *m/z* range for MS^2^ scan pairing (Da): 0.01; RT range for MS^2^ scan pairing (min): 0.2; (iv) Isotopic peaks grouper: *m/z* tolerance: 0.001 *m/z* or 15ppm; Retention time tolerance: 0.2; Maximum charge: 3; Representative isotopes: Most intense; (v) Join aligner: *m/z* tolerance: 0.001*m/z* or 15ppm; Weight for *m/z*: 75; Retention time tolerance: 0.2; Weight for RT: 25; (v) Peak finder: Intensity tolerance: 10; *m/z* tolerance: 0.001*m/z* or 15ppm; Retentation time tolerance: 0.2; (vi) Export to CSV file: Export common elements: Export row ID, Export row *m/z*, Export row retention time; Export data file elements: Peak area; Peak *m/z* min; Peak *m/z* max; Filter rows: All.

The following parameters have been adopted for the MS/MS data acquired in DDA mode: (i) Mass detection (MS level 1): mass detector – Centroid; noise level – 1E3; (ii) Mass detection (MS level 2): mass detector – Centroid; noise level – 1E2; (iii) ADAP: min group size – 4; Group intensity threshold – 1E3; Min highest intensity – 3E3; *m/z* tolerance – 0.001*m/z* or 15ppm; (iv) Chromatogram deconvolution: algorithm – Baseline cut-off; Min peak height – 3E3; Peak duration range: 0 – 15; Baseline level – 1E3; *m/z* center calculation: Median; *m/z* range for MS^2^ scan pairing (Da): 0.01; RT range for MS^2^ scan pairing (min): 0.2; (v) Isotopic peaks grouper: *m/z* tolerance: 0.001*m/z* or 15ppm; Retention time tolerance: 0.2; Maximum charge: 3; Representative isotopes: Most intense; (vi) Join aligner: *m/z* tolerance: 0.001*m/z* or 15ppm; Weight for *m/z*: 75; Retention time tolerance: 0.2; Weight for RT: 25; (vii) Peak finder: Intensity tolerance: 10; *m/z* tolerance: 0.001*m/z* or 15ppm; Retention time tolerance: 0.2; (viii) Export to CSV file: Export common elements: Export row ID, Export row *m/z*, Export row retention time; Export data file elements: Peak area; Peak *m/z* min; Peak *m/z* max; Filter rows: All; (ix) Export/Submit to GNPS-FBMN: Filter rows: Only with MS^2^.

The following parameters have been adopted for the MS/MS mode of data obtained with the scheduled precursor list (SPL): (i) Mass detection (MS level 1): mass detector – Centroid; noise level – 1E2; (ii) Mass detection (MS level 2): mass detector – Centroid; noise level – 1E1; (iii) ADAP: min group size – 4; Group intensity threshould – 1E2; Min highest intensity – 3E2; *m/z* tolerance – 0.001*m/z* or 15ppm; (iv) Chromatogram deconvolution: algorithm – Baseline cut-off; Min peak height – 3E2; Peak duration range: 0 – 15; Baseline level – 1E2; *m/z* center calculation: Median; *m/z* range for MS^2^ scan pairing (Da): 0.01; RT range for MS^2^ scan pairing (min): 0.2; (v) Isotopic peaks grouper: *m/z* tolerance: 0.001*m/z* or 15ppm; Retentation time tolerance: 0.2; Maximum charge: 3; Representative isotopes: Most intense; (vi) Join aligner: *m/z* tolerance: 0.001*m/z* or 15ppm; Weight for *m/z*: 75; Retentation time tolerance: 0.2; Weight for RT: 25; (vii) Peak finder: Intensity tolerance: 10; *m/z* tolerance: 0.001*m/z* or 15ppm; Retentation time tolerance: 0.2; (viii) Export to CSV file: Export common elements: Export row ID, Export row *m/z*, Export row retention time; Export data file elements: Peak area; Peak *m/z* min; Peak *m/z* max; Filter rows: All; (ix) Export/Submit to GNPS-FBMN: Filter rows: Only with MS^2^.

In all cases, a batch mode file was made available in the Zenodo repository in the Data Availability session.

#### 2.2.2 Ion filtering and schedule precursor list (SPL) construction

RegFilter is an open-source software written in Python and available from the laboratory’s GitHub repository (https://github.com/computational-chemical-biology/regression_filter). To use it, the user must follow the installation steps described in the repository. Once installed, RegFilter uses as input material a table in .csv format containing the spectral data previously acquired by the user in full scan mode and duly pre-processed by MZmine (Schmid et al., 2023) software as described above. The package’s instructions guide user on how to use the package and export a report showing the number of total features, the number of filtered features, and a new .csv table with the filtered features with their respective retention times, that can be formatted as a SPL for different mass spectrometers.

#### 2.2.3 GNPS2 Molecular Networking and *in silico* annotation

From the raw data of the spectral analysis, we preprocessed these data through MZmine (Schmid et al., 2023), generating an area below the chromatographic curve quantification table (.csv) and a list of fragmentation spectra (.mgf) files which were imported into the GNPS (Wang et al., 2016) and GNPS2 platforms (gnps2.org/), where the molecular network and spectral pairing annotation was performed by the Feature Based Molecular Network (FBMN) workflow (Nothias et al., 2020). For the nodes that could not be annotated by spectral library matching, the MS/MS mass spectra were annotated in two layers of annotation. First, we propagated annotations based on the topology of the molecular network using the ChemWalker tool (Borelli et al., 2023) through its new graphical interface available on GNPS2 (https://gnps2.org/workflowinput?workflowname=chemwalker_nextflow_workflow).

The criteria used to propagate the annotation were: (i) Adduct: [M+H]^+^; (ii) tolerance: 15 ppm. A second layer of *in silico* annotation was then performed using the SIRIUS tool (Dührkop et al., 2019) version 5.8.5 for the nodes that could not be annotated in the previous steps. The recommended parameters for the Q-TOF instrument were adopted, using the formulas available in all databases and taking into account the adducts [M+H]^+^, [M+K]^+^, and [M+Na]^+^. In both cases of *in silico* annotation, we considered the best-ranked candidate as the annotation for the corresponding spectrum.

## 3 Results and Discussion

### 3.1 Regression Filter: optimizing the detection of biological signals through a semi-automated feature selection tool

Some of the bottlenecks that deserve to be highlighted in the processing and analysis of metabolomics data from untargeted experiments are (i) the selection of features directly related to the sample, i.e., biological signals, and (ii) the fragmentation of features that are lost in DDA. To overcome these limitations, we have created a dynamic analysis flow that combines experimental and computational approaches, as illustrated in Figure 1. In this analysis flow, a sample of interest is diluted serially (until 8-fold dilution). An LC-MS analysis is performed on the undiluted sample and each dilution in full scan mode. The raw data are converted to the .mzXML format and then pre-processed by the MZmine software (Schmid et al., 2023) with the parameters that best fit the user’s necessities. The .csv file obtained in this step is subjected to feature filtering using an algorithm called Regression Filter (RegFilter). This algorithm is implemented in Python and describes the conditions to filter features. All the ions presented in the entire dilution series whose area under their curve decreases linearly, respecting a coefficient of determination (R^2^) greater than or equal to 0.9 (default value that can be adjusted by the user) and a negative angular coefficient for the fitted regression model, are stored and reported to the user. The output is then exported as a new .csv file containing all the selected features with their respective retention times, which is then inserted into the mass spectrometer software as a scheduled precursor list (SPL), where the ions it contains are fragmented only in the undiluted sample.

**Figure 1.**
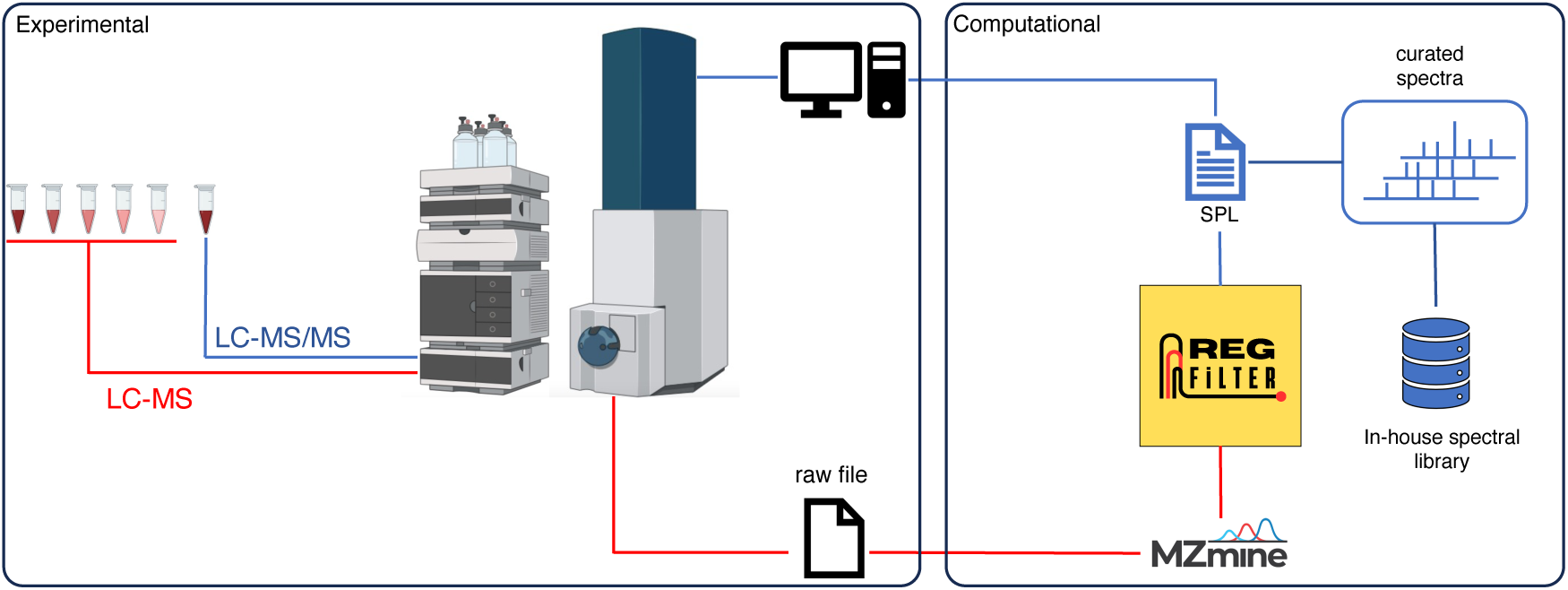
Analysis workflow of an untargeted metabolomics experiment using the RegFilter platform.

In addition to the feature filtering algorithm itself, the package provides several functions for data analysis and visualization as well as the possibility of storing spectra of interest for *in-house* library building. To test the tool’s efficiency in extracting features of potential biological relevance, as well as its potential to select features previously lost in the DDA mode, we performed a series of experiments in which the cellular extract of three marine actinomycetes (BRA006, BRA010, and BRA177) belonging to a Brazilian collection of bacterial isolates was subjected to the workflow, and the results obtained were compared with a dataset of fragmentation spectra obtained in DDA, the most common acquisition mode for untargeted experiments in complex biological samples. The choice to work on this biological matrix is due to the potential for the production of bioactive compounds that this group of bacteria represents, opening the possibility for the discovery, cataloging, and curation of new chemical entities (van Bergeijk et al., 2020; Chen et al., 2021).

### 3.2 RegFilter efficiency on chromatographic run coverage

We analyzed whether RegFilter was able to cover the entire chromatographic run. As can be seen in Figure 2 with the chromatograms 3D plot where we compared SPL and DDA chromatograms from those ions that led to fragmentation, RegFilter covered the entire analysis region from the three bacterial extracts. It is worth noting that ions with a low mass/charge ratio (*m/z*) were more frequently captured by the list compared to DDA, where the prevalence of these precursors destined for fragmentation throughout the chromatographic run is clear. Additionally, a less noisy baseline is observed in these chromatographic runs in all extracts, indicating that the noise filtering process was efficient.

**Figure 2.**
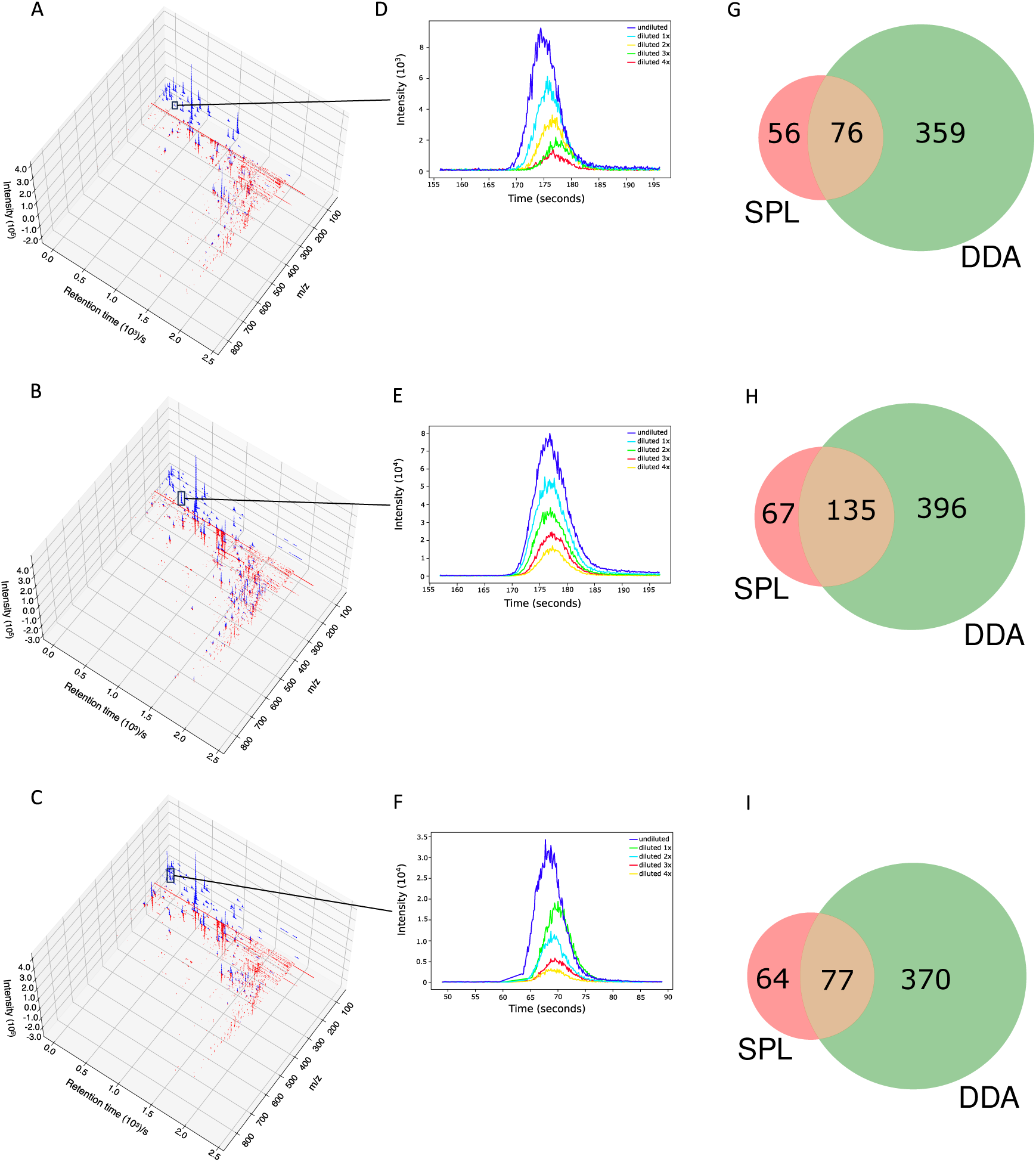
RegFilter was able to efficiently cover all the chromatographic runs. (A-C) 3D chromatograms for the fragmented precursor ions for the three actinomycetes analyzed, where (A) BRA006, (B) BRA010, and (C) BRA177. The blue and red lines indicate the precursor ions selected for fragmentation with their respective intensities selected by SPL and DDA, respectively. (E-F) Extracted ion chromatograms (EIC) of meaningful examples of real signals selected by RegFilter. (G-I) Venn diagrams showing the overlap between the precursor ions subjected to fragmentation by DDA mode and scheduled precursor list (SPL) generated through RegFilter.

Once verified that RegFilter could cover the entire analysis region, we proceeded with a first comparative analysis between the full scan and data-dependent acquisition (DDA) modes. The results are shown in Table 1. We can see that the total number of features detected in full scan mode is higher than the total number of features detected in DDA mode. This is an expected event because the instrument spends a considerable amount of time generating the spectra at the MS^2^ level, which reduces the intensity of the features at the MS^1^ level and, consequently, their detection, meaning that some of the ions previously detected in full scan mode are no longer detected (Guo and Huan, 2020).

**Table 1.**
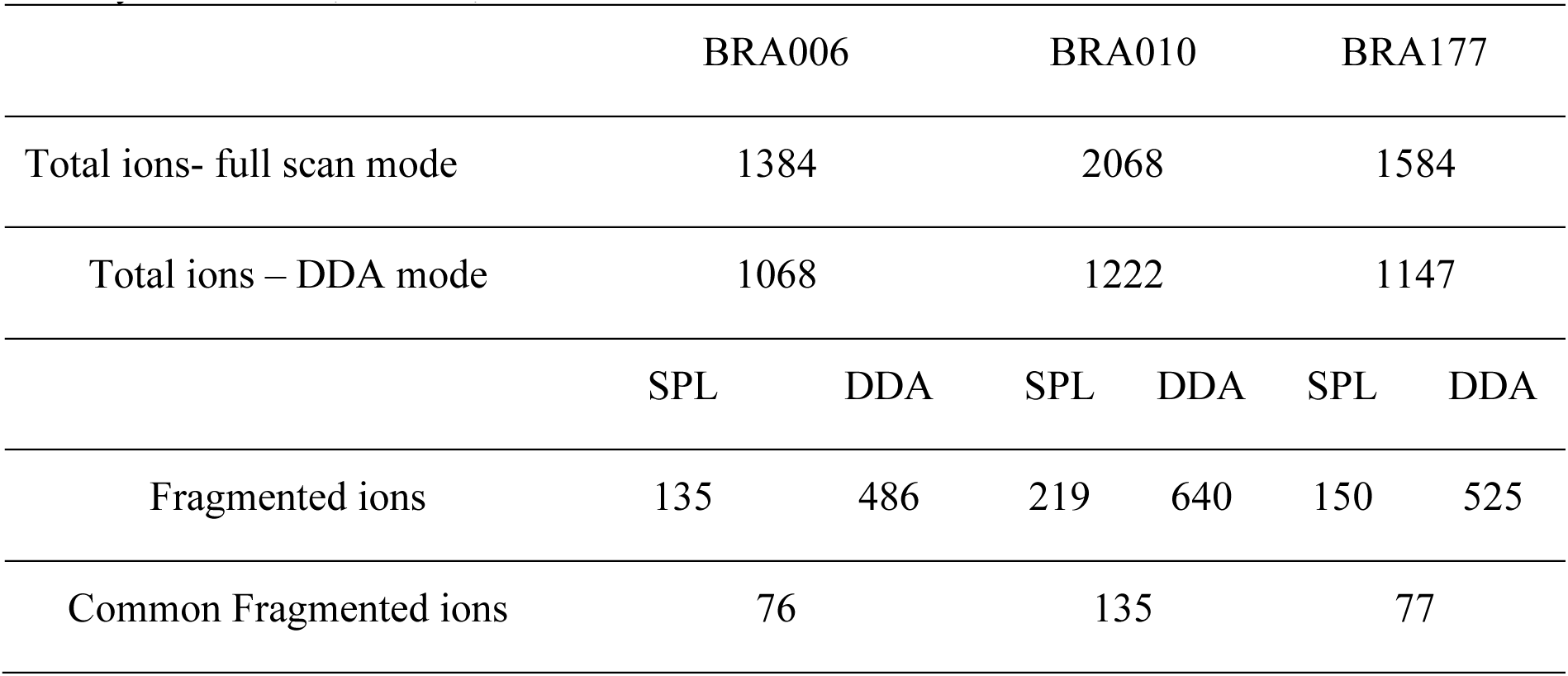
Comparison between data obtained in untargeted metabolomics experiments for each of the actinomycetes BRA006, BRA010, and BRA177.

|  | BRA006 |  | BRA010 |  | BRA177 |  |
| --- | --- | --- | --- | --- | --- | --- |
| Total ions- full scan mode | 1384 |  | 2068 |  | 1584 |  |
| Total ions – DDA mode | 1068 |  | 1222 |  | 1147 |  |
|  | SPL | DDA | SPL | DDA | SPL | DDA |
| Fragmented ions | 135 | 486 | 219 | 640 | 150 | 525 |
| Common Fragmented ions | 76 |  | 135 |  | 77 |  |

Applying RegFilter to the features detected in full scan mode, considering a regression fitted with at least three points (the experiments were conducted with 5 experimental points), it can be seen that 135 features for strain BRA006, 219 features for strain BRA010, and 150 features for strain BRA177 met the filtering requirements. Based on this result, we wondered how many and which features would be common to those detected by the DDA mode. As shown in the Venn diagrams (Figure 2 G-I) and Table 1, we observed that 76 features, 135 features, and 77 features for strains BRA006, BRA010, and BRA177, respectively, were present both in the scheduled precursors list (SPL) selected by RegFilter and in the fragmented precursor ions in DDA mode.

It is worth noticing that there is a difference between the number of fragmented features shown in Table 1 and Figure 2G-I, both in the list and the DDA. Even with the pre-processing approach performed through MZmine, there may be redundancy in the detected features, which is common in data obtained in metabolomics experiments (Zhang et al., 2023). Therefore, the observed difference is due to the reduced redundancy verified by creating a unique identifier that combines the *m/z* values and retention times for each feature. Such diminished redundancy was achieved through a mass and retention time matching function contained in the RegFilter package. From these common precursor ions, we evaluate their fragmentation profile, taking as a basis for comparison (i) the number of peaks generated by fragmentation, (ii) the average, (iii) the maximum and (iv) minimum intensities of these peak heights, and of (v) the number of events in MS^2^, which is how many times a given precursor ion was directed to fragmentation. Table 2 shows the number of precursor ions, with a higher average for those ions presented in three replicates (“upregulated”), showing statistically significant differences in the parameters described above (p<0.05, Student’s t-test).

**Table 2.** Comparison between parameters of the fragmentation spectra between SPL and DDA modes (p<0.05, Student’s t-test).

|  | BRA006 |  | BRA010 |  | BRA177 |  |
| --- | --- | --- | --- | --- | --- | --- |
|  | SPL | DDA | SPL | DDA | SPL | DDA |
| Number of peaks | 5 | 22 | 4 | 55 | 5 | 20 |
| Mean intensity | 1 | 26 | 6 | 53 | 3 | 40 |
| Maximum intensity | 4 | 28 | 4 | 64 | 6 | 34 |
| Minimum intensity | 3 | 4 | 25 | 5 | 5 | 2 |
| MS<sup>2</sup> events | 54 | 1 | 114 | 0 | 64 | 8 |
| Percentage of noisy spectra* | 4,4 | 4,9 | 5,9 | 3,8 | 4,7 | 4,4 |

\*Manually inspected

Of those ions that showed a statistically significant difference in the mean, except the number of events in MS^2^ for all strains and the minimum intensity for strains BRA010 and BRA177, the data-dependent acquisition (DDA) mode showed a better performance in obtaining fragmentation spectra compared to fragmentation spectra generated by list targeting. The quality of fragmentation spectra can be measured by evaluating the number of fragments (Ausloos et al., 1999; Bern et al., 2004; Zhang et al., 2023) and the signal intensity of the detected features (Bern et al., 2004; Broeckling et al., 2018). Even with such quantitative prerogatives, the proper assessment of the quality of a given spectrum depends critically on the proper judgment and expertise of a spectrometrist, as well as the pairing of the spectrum of a given compound of interest with a reference spectrum (Ausloos et al., 1999).

Intuitively, since the number of events in MS^2^ for SPL was higher for all strains compared to the number of events in MS^2^ for DDA, we expected that the quality of the fragmentation spectra would, consequently, be higher for the fragmented ions in SPL than in DDA, which was not verified. However, it is important to highlight that not all ions common between SPL and DDA in the three bacterial strains showed a statistically significant difference in the parameters evaluated. In fact, for most of these common ions, except the events in MS^2^, the spectral comparison based on the evaluated parameters did not show a statistically significant difference. Since the entire basis of the spectral comparison is based on the spectra of interest against reference spectra, we decided to go one step further in comparing these spectra. Therefore, we took the common ions fragmented by both the list and DDA and performed their spectral pairing separately in the GNPS library (Wang et al., 2016). For the spectra that were annotated and had the same annotation for the same precursor ion present in both the SPL and DDA, we evaluated the number of common peaks relative to the reference spectra. The results of this comparison are shown in Supplementary Figure 1.

Comparing the average number of common peaks of the same annotated ions between SPL and DDA, there is no significant statistical difference between them. Therefore, the quality of the fragmentation spectra generated by SPL and DDA has the same annotation potential. Taking into account the context of the analysis, the collision energy used was the same for the ions contained in both the SPL and the DDA, which helps to explain this absence of difference observed. Next, we decided to test the hypothesis that the use of fragmentation lists would lead to a selection of less noisy features. For this purpose, we used the construction of extracted ion chromatograms (EIC) for each of the precursor ions included in both the list generated by RegFilter and the precursor ions fragmented by DDA for each of the three model strains studied. The distinction between signal and noise is given by the shape of the curve of the extracted ion chromatogram. A true signal has a well-defined sinusoidal shape, whereas the noise is characterized by its random distribution profile (Jankevics et al., 2012).

Using this profile as an evaluation reference, we manually inspected and analyzed each of the chromatograms, and quantified the percentage of noise for all three strains, both from the precursors included in the fragmentation list (SPL) and DDA mode. These results are presented in Table 2 (percentage of noisy spectra). It can be seen that there is no real difference in the amount of noise between the ions generated from the list (SPL) and those from the DDA. Detecting noisy features is an inherent challenge in MS-based metabolomics experiments (Beisken et al., 2015). One of the pitfalls of analyses performed with this analytical technique, especially when considering time-of-flight analyzers, is that signals of reproducible intensities tend to be generated in noisy regions (Sands et al., 2021). In this sense, the practice of serial dilution is widely used as a resource to identify such noisy features, allowing the selection of quality features and acting as more reliable and interpretable data (Jankevis et al., 2012; Sands et al., 2021). The reproducibility of such noisy signals described by Sands and co-workers (2021) was confirmed in our results, as even with a five-point regression model, these noisy signals persisted throughout the dilution curve. In other words, the filtering process based on a linear regression model alone does not guarantee the exclusion of noisy features, requiring an additional selection step. In this sense, RegFilter proved to be an excellent resource in the process of presenting, detecting, and selecting quality features, as it allows the user to visualize and discriminate what these features are.

It is important to note that the reduction of noise and better-quality fragmentation spectra has been previously reported in the literature (Jankevics et al., 2012, Broeckling et al., 2018). The fact that these observations were not replicated in the present study is partly due to the low complexity of the bacteria extract used. More complex matrices, such as plants or environmental samples, may present higher complexity and benefit from noise removal and increased spectral quality in addition to the complementary coverage demonstrated here.

### 3.3 3.3 Flying Under the Radar: Detecting once-missed signals in data-dependent acquisition mode and exploring the potential of in-house MS libraries

Turning our attention to the features contained only in SPL, we decided to look for annotations for precursor ions, and their respective fragmentation spectra, contained only in the list. To investigate the consistency of the features, we further inspected the presence of the precursor ions (*m/z*) and the corresponding retention times for each of the precursor ions present in the list in the DDA MS^1^ data, i.e. the total features. The overlap between the precursor ions in the list and the total features detected in DDA was 99.2%, 99.0%, and 100.0% for strains BRA006, BRA010, and BRA177, respectively (Figure 3). In other words, almost all of the precursor ions selected in the fragmentation list were present in the MS^1^ of the DDA method, but not all were fragmented. We decided to go one step further in analyzing this result. Therefore, we manually examined each of these ions that were exclusively fragmented by the list and were present in MS^1^ of the DDA, but not fragmented in the latter mode. We found that in 33.9% of cases for strain BRA006, 22.4% of cases for strain BRA010 and 29.7% of cases for strain BRA177 the base peak (most intense ion on the MS^1^ spectrum) was the ion fragmented by the list, and not fragmented by DDA. For the other ions fragmented by the list, the peaks selected were not the base peak, having intensities from 1.1 to 86.2 times lower than the highest intensity ion for BRA006, from 1.2 to 46.5 for BRA010, and from 1.2 to 57.1 for BRA177, which explains their lack of fragmentation by the data-dependent acquisition mode. One of the limitations of the data-dependent fragmentation method is precisely its stochastic nature, where ions with similar intensities may or may not be fragmented (Davies et al., 2021). This finding was only possible by manual verification and analysis of these data. Even in the case of the smallest fraction of fragmented ions included in the list, it is still noteworthy because even those ions that would theoretically be good candidates for fragmentation by the DDA mode were not fragmented and, most importantly, RegFilter was able to select these ions.

**Figure 3.**
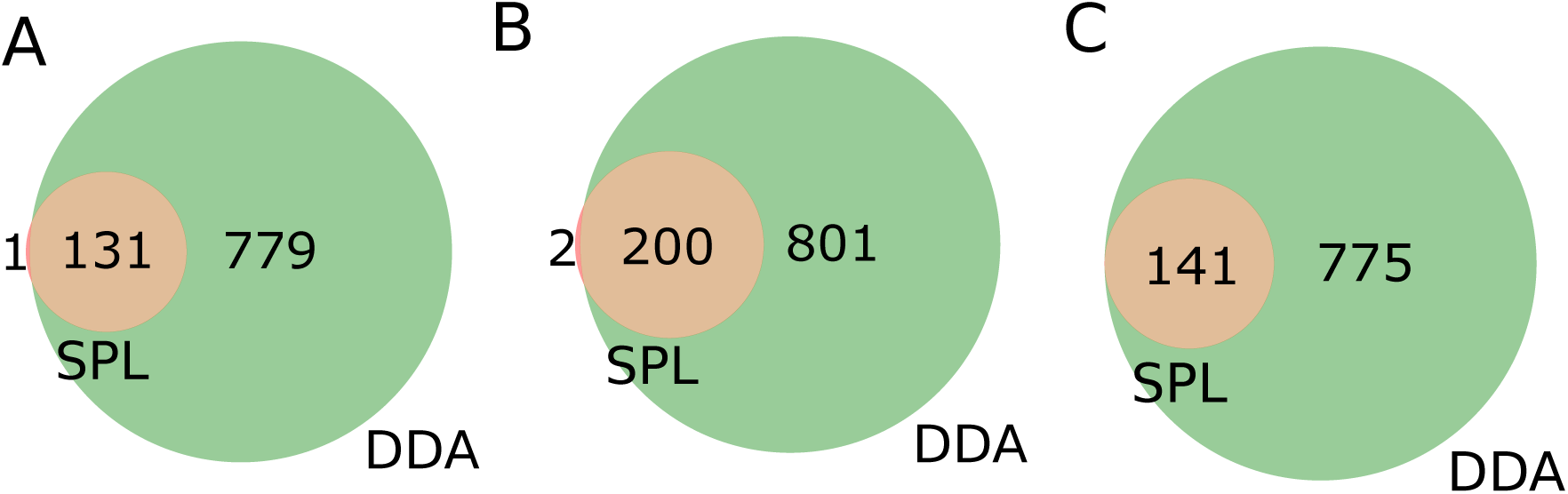
Venn diagrams of features present in the fragmentation list (SPL) and detected among the total features contained in DDA mode. (A) BRA006, (B) BRA010 and (C) BRA177.

Now, to look for annotations in the fragmentation spectra of the ions selected from the list, we took three approaches: first, we performed their spectral pairing using the GNPS2 library (Wang et al., 2016), as it comes from one of the largest public libraries of spectral data (Bittremieux et al., 2022). For those features that could not be annotated using this approach, we decided to go one step further and use ChemWalker to propagate the annotations through the topology of the molecular network (Borelli et al., 2023). Finally, we used the SIRIUS tool (Dührkop et al., 2019) on the fragmentation spectra of each of the candidates that could not be annotated through GNPS spectral pairing as well as ChemWalker annotation propagation. It should be noted that none of the annotated spectra was considered noise, but rather a true signal. As a result, 43 annotations were recovered for BRA006, 46 annotations for BRA010, and 47 annotations for BRA177. All the annotations are presented in the Supplementary Table 1.

To gain further information on the biological interpretation of the identified molecules, we cross-referenced the annotation data of each of the molecules contained in the three bacteria with the KEGG database (Kanehisa and Goto, 2000) to assess whether the set of molecules detected had already been described and cataloged in the encyclopedia along with its metabolic pathway presented. The compounds annotated with their respective pathway are presented in Supplementary Table 2. From the total set, 39 annotated compounds were identified using KEGG with their respective metabolic pathway associated. Of which, nine for BRA006, 10 for BRA010, and 20 for BRA177 were described along known pathways. The relationship between the compounds described and their respective pathways is shown in Supplementary Table 2. We can see that there are compounds common to the three strains, such as styrene, and others, such as 2-methylbenzaldehyde, that are common to BRA006 and BRA010. In turn, it is interesting to note that there are common metabolic pathways between the three strains but with different intermediates, such as purine metabolic pathways. Particularly in the case of the BRA177 compound, it is interesting to note that of the 20 annotated features, three had the same annotation for three different features, indicating the possible presence of isomers and/or adducts. However, the most important aspect of this result is that list-directed feature fragmentation makes it possible to annotate biological signals previously lost in DDA.

As mentioned in the tool description, with the RegFilter package, it is possible to build in-house libraries to store spectra of interest using a html-based graphical interface included in the package. The sequence of steps to create such a library is shown in Supplementary Figure 2. We created a small library by focusing our attention on the ions targeted for fragmentation by the list, and with this repository in hand, we followed two different approaches, as shown in Figure 1. First, using the MASST tool (Wang et al., 2020), we searched the individual spectra against public spectrometric data, which greatly expands the potential of public spectral libraries. Of those that matched the platform, the spectrum with the highest score was downloaded, and, through a local search function, that is part of the package, we performed the reverse path. In other words, we show how one could take a spectrum obtained from a public database, perform spectral matching, calculate the similarity score, and evaluate the number of peaks shared between this reference spectrum and all those contained in one’s local library. Figure 4A-C shows the results of examples to showcase the potential of local spectral collections, where the cosine score values and number of paired peaks for actinomycetes BRA006 (6A), BRA010 (6B), and BRA177 (6C) were 0.9565 and 25, 0.3228 and 10, 0.6048 and 13, respectively. In addition, we analyzed the spectra contained in the in-house libraries for the three strains with microbeMASST (Zuffa et al., 2024) to evaluate the potential of these spectra to provide evidence for the taxonomy of these actinomycetes through this selected list of spectra. It was possible to find similarities between them and other spectra described for other microorganisms (Figure 4D). These results illustrate the potential and advantages of building custom libraries to obtain chemical information. It also demonstrates the potential of the Regfilter tool for obtaining spectra of relevant biological importance.

**Figure 4.**
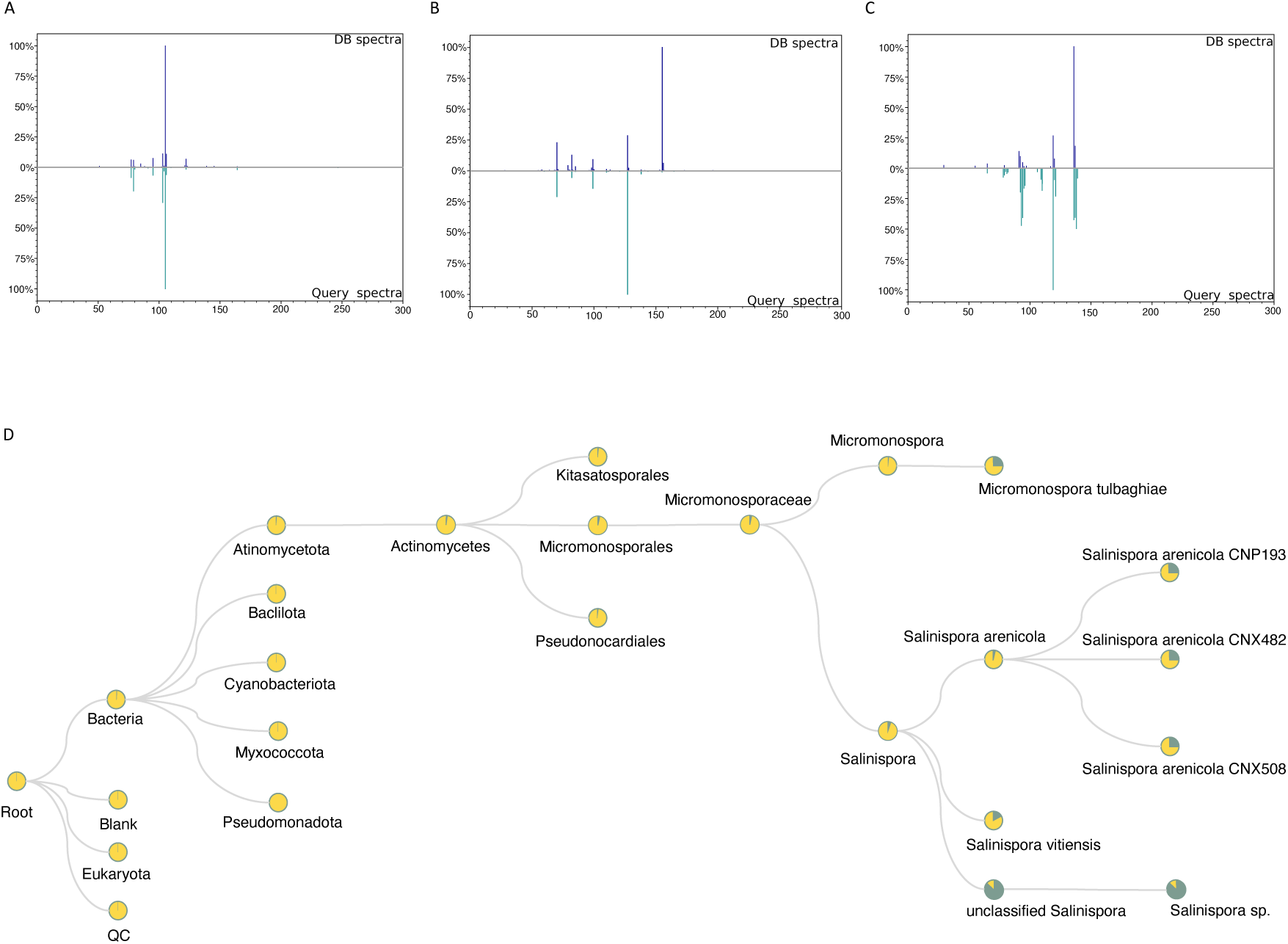
The spectra contained only the chemically and biologically relevant information. 4A-C: Examples of mirror plots between the best matches among reference spectra extracted from GNPS and the in-house database, for the actinomycetes BRA006 (A), BRA010 (B), and BRA177 (C). (D) Representative cladogram indicating the interspecies proximity of a given metabolite that is exclusively included in the list (SPL).

Among the current challenges in the design of experiments and tools that allow both the acquisition and the efficient use of metabolomics data based on mass spectrometry, the selection of features relevant to a context of interest, allowing a targeted experiment after an untargeted, is one of them. In this sense, the results obtained by the hybrid approach proposed in this work, with emphasis on the RegFilter package, have shown that this tool is not only capable of selecting biologically relevant features but also allows the fragmentation of features previously lost by the DDA mode. Proposals for the use of serial dilution in mass spectrometry-based metabolomics experiments have been suggested in the literature with different goals (Jankevics et al., 2012; Duan et al, 2016; Sands et al., 2021; Zhang et al., 2023). The focus of Duan and co-workers (2016) was to overcome the problem of noise inherent in experiments using mass spectrometry as an analytical resource. To overcome this problem, the authors used serially diluted quality control samples to detect signals that show a linear pattern of quantification in the spectrometry and are, therefore, not due to random noise. The authors reported that the method was able to reduce signals detected from a mixture of 20 known compounds, with 1,342 mass peaks detected. After filtering, only 102 mass peaks (7.6%) were recovered and assigned to the 20 original compounds. When considering our results, RegFilter was able to select and recover 9.3% of BRA006, 9.9% of BRA010, and 9.0% of BRA177 biological signals from the total number of features observed in MS^1^ full scan mode. Despite the difference in recovering signals from standard samples, these numbers align with the finding that mass spectra are highly redundant and noisy. Still, in terms of recovering potentially relevant signals, this is a considerable recovery since they are true signals and can be considered of higher quality and reliability (Sands et al., 2021). It should be noted that the process was carried out in a semi-automated way, and these are relatively low complexity samples if compared with plant or environmental extracts, where more signal and more noise are expected. The results highlight the potential for applying linear regression methods for more complex samples where the proportion of signal/noise is smaller.

RegFilter was able to select ions of low relative abundance, overcoming one of the limitations already described for this acquisition mode itself (Zhang et al., 2023). Another important difference in using this analytical tool is that RegFilter proved to be effective in what could be called the enrichment of data with potential biological value since the features detected by it proved to be stable throughout the dilution steps and, therefore, exclusively related to the biological sample. Several works point to the need to increase the coverage of ions to be fragmented in untargeted metabolomics experiments. Multiple injections of the same sample (Broeckling et al., 2018), selecting the totality of ions contained in specific regions of the chromatogram (Davies et al., 2021), and automating the exclusion of ions to avoid redundancy (McBride et al., 2023) are some of the many strategies to increase the number of ions to be fragmented in a given context of interest. However, in addition to increasing the fragmentation coverage, as important as the number of fragmented ions is the choice of which of these ions are targeted for fragmentation. In this sense, the debate on the fragmentation of specific ions mediated by inclusion lists has been a recent topic (Koelmel et al., 2017; Broeckling et al., 2018; Zhang et al., 2023). In the case of economically expensive experiments (Aksenov et al., 2017), a more targeted selection would allow the optimization of time and resources in discovering molecules of biological interest.

Recently, Zhang and co-workers (2023) presented improvements and advantages of using a list of pre-selected precursors to gain biological information in untargeted metabolomics experiments. To this end, the list of precursors is constructed by analyzing the samples of interest in full scan mode, and based on these data and which ions would be statistically different in MS^1^ full scan mode between contrasting experimental groups of samples, the list is constructed, and the ions contained therein are fragmented. The approach presented by the authors has important parallels with our approach in RegFilter, where both first use MS^1^ full scan data to construct the inclusion lists. However, the inclusion criteria are different and this is where RegFilter stands out for the novelty of its approach. Firstly, the selection of features is semi-automatic, allowing full automation, as the software can be applied to a serial dilution, and the list generated can be communicated to the instrument to fragment samples that are obtained in sequence (Broeckling et al., 2018), which itself represents an optimization in the time for data acquisition and analysis. The feature selection criteria promoted by RegFilter are essentially based on two aspects: provided that the features contained in a dilution series are consistent with the construction of a curve with a coefficient of determination (R2) greater than or equal to 0.9 and with a negative slope coefficient, these features are included in the fragmentation list. Zhang and coworkers (2023), on the other hand, used the construction of lists with control samples to extrapolate the data to real samples. In this sense, RegFilter can build lists from the samples themselves simply by diluting them.

These different approaches can be complementary, as the filter points out noisy features that may not be removed by a differentially regulated approach. The main drawbacks of serial dilutions are additional data collection and features that have low intensity and may not present in at least three less diluted samples to fit a curve. Of all the challenges in mass spectrometry-based metabolomics, one of the greatest is undoubtedly the design and execution of experiments that allow for more effective targeting when collecting and navigating the sea of data generated in this field. Even with increasing access to cutting-edge techniques, improved data analysis capabilities, and advances in technology to build thinner and more accurate spectrometers, a lot of information is lost in untargeted analysis. Or worse, a lot of data is discarded, and data replication using the untargeted approach on the same biological matrix is performed repeatedly. In this respect, RegFilter is a promising tool, not only to guide the acquisition of data relevant to the sample of interest but especially to allow the acquisition and selection of data to be curated and stored, allowing the construction of spectral libraries of specific interest.

## 4 Conclusions

In conclusion, the adoption of an analysis flow that dynamically integrates experimental and computational approaches, with the latter introducing the RegFilter feature selection tool, has made it possible to optimize the detection of biological signals belonging to the sample of interest that were previously lost in the traditional Data-Dependent Fragmentation (DDA) mode. Furthermore, RegFilter package opens up the possibility of creating curated spectral libraries with relevant biological information.

## 5 Author contributions

GSA: conceptualization, data curation, formal analysis, investigation, methodology, software, validation, visualization, writing – original draft, writing – review and editing. LGM: data curation, methodology, software, validation, writing – review and editing. RdF: data curation, investigation, validation, writing – review and editing. LGPF: data curation, validation, writing – review and editing. PR: data curation, writing – review and editing. HT: data curation, software, writing – review and editing. AP: data curation, writing – review and editing. DRP: data curation, writing – review and editing. LVCL: funding acquisition, project administration, resources, writing – review and editing. NPL: funding acquisition, project administration, supervision, writing – review and editing. DBBT: funding acquisition, project administration, resources, writing – review and editing. RRdS: conceptualization, data curation, formal analysis, investigation, methodology, software, validation, visualization, funding acquisition, project administration, supervision, writing – original draft, writing – review and editing.

## 7 Acknowledgments

Sample collection and bioprospection authorizations were granted by the Biodiversity Authorization and Information System (SISBIO), number 26286-1 for the collection of UFC, from which BRA-006, BRA-010 and BRA-177 were isolated

## 9 Data Availability

All mass spectrometry metabolomics data are available at MassIVE through the identifiers: MSV000095042, MSV000095044, and MSV000095129. RegFilter package can be found at https://github.com/computational-chemical-biology/regression_filter. The parameters used for preprocessing in MZmine are available at Zenodo https://doi.org/10.5281/zenodo.10822821 as .xml forms. The links for the jobs made on GNPS2 are available on: https://gnps2.org/status?task=834d956aa8b54757915e1a93af2866b4; https://gnps2.org/status?task=b7da5a5d55334f3090d07712971df236; https://gnps2.org/status?task=1abad20ab0b44181a0421a4371b5ca00; https://gnps2.org/status?task=96c2d2be678a45718050fe4aca5a5bfc; https://gnps2.org/status?task=f1e2242d9aad4fca9d5bec3746fae479; https://gnps2.org/status?task=d6fd31a18cc942739096b7cf7f338cde

The links for the jobs made on GNPS are available at: https://gnps.ucsd.edu/ProteoSAFe/status.jsp?task=51abc1f1bfbe4a4fa3da3ace237f90d0; https://gnps.ucsd.edu/ProteoSAFe/status.jsp?task=e39178dcf8f142649a00f425f9a52b47; https://gnps.ucsd.edu/ProteoSAFe/status.jsp?task=8d5285bb93e44c149ba09c9a86930f53

