## Supplemental material for "Optimizing the detection of biological signals through a semi-automated feature selection tool"

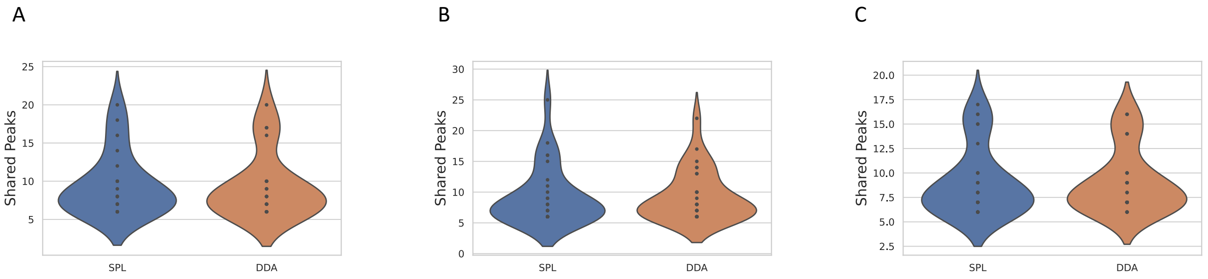


Supplementary Figure 1. Violin plots comparing the average peaks shared by the same precursor ions in SPL and DDA when paired with reference spectra in the GNPS2 library (Wang et al., 2016), namely (A) BRA006, (B) BRA010 and (C) BRA177.

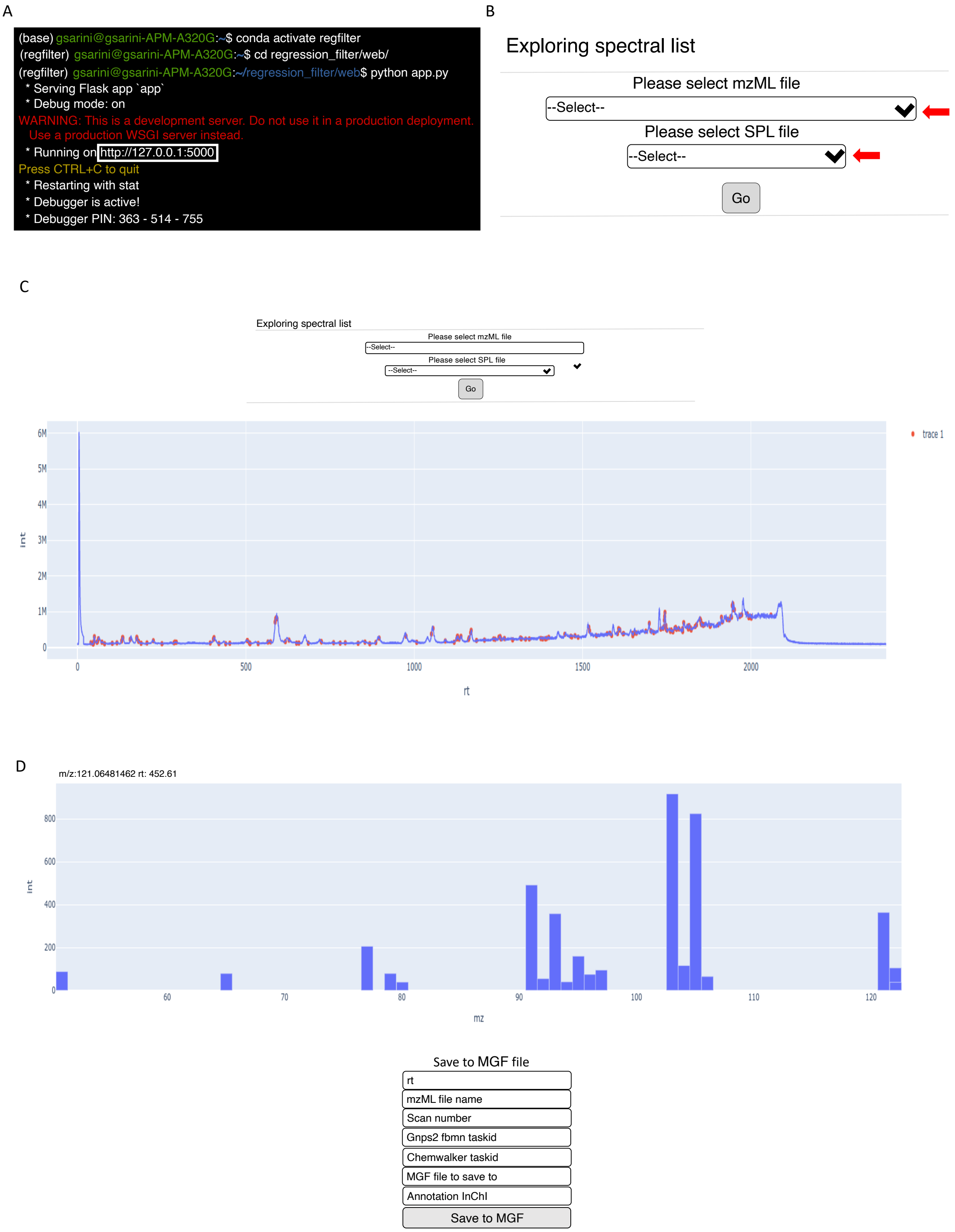


Supplementary Figure 2. Step-by-step instructions for creating in-house spectral libraries using RegFilter. After installing RegFilter follow the additional graphical interface instructions (https://github.com/computational-chemical-biology/regression_filter/tree/main/web). By clicking on the link shown in the white box (A), the user is taken to their default web browser. There (B), the user selects the spectrometric data of interest and the desired fragmentation list. Once both have been selected, the spectrum is displayed, with the red dots in the fragmentation list (C) indicating the features included in the list. By clicking on one of the points of interest, the user is presented with the corresponding fragmentation spectrum and can make insertions of interest in the .mgf file to be exported (D).

Supplementary Table 1. Molecular annotations obtained from the GNPS, SIRIUS and ChemWalker tools for the features included only in the list for Actinomycetes BRA006, BRA010, and BRA177.

| Sample | Precursor Mass (m/z) | Retention time (min) | Annotated Method | InChIKey | IUPAC name |
|---|---|---|---|---|---|
| BRA006 | 136.076 | 0.05 | ChemWalker | ZXTLGJAARBNQGK-UHFFFAOYSA-N | N-(2-methylphenyl)formamide |
| BRA006 | 108.045 | 0.81 | ChemWalker | CSDSSGBPEUDDEE-UHFFFAOYSA-N | pyridine-2-carbaldehyde |
| BRA006 | 96.045 | 0.82 | ChemWalker | ZSKGQVFRTSEPJT-UHFFFAOYSA-N | 1H-pyrrole-2-carbaldehyde |
| BRA006 | 126.055 | 0.83 | ChemWalker | VJBRLHBYLMMWER-UHFFFAOYSA-N | N-[(5-methylfuran-2-yl)methylidene]hydroxylamine |
| BRA006 | 110.060 | 0.83 | ChemWalker | OUKQTRFCDKSEPL-UHFFFAOYSA-N | 1-methylpyrrole-2-carbaldehyde |
| BRA006 | 167.093 | 0.84 | SIRIUS | PVWNFAGYFUUDRC-UHFFFAOYSA-N | N-[(4-amino-2-methylpyrimidin-5-yl)methyl]formamide |
| BRA006 | 123.047 | 1.09 | ChemWalker | YODZTKMDCQEPHD-UHFFFAOYSA-N | 2-(2-hydroxyethylsulfanyl)ethanol |

| | | | | | |
|---|---|---|---|---|---|
| BRA006 | 137.071 | 1.37 | ChemWalker | RHPIFAMEPPCHGX-UHFFFAOYSA-N | 6-(hydrazinylmethylidene)cyclohexa-2,4-dien-1-one |
| BRA006 | 153.041 | 1.57 | GNPS2 | LRFVTYWOQMYALW-UHFFFAOYSA-N | 3,7-dihydropurine-2,6-dione |
| BRA006 | 174.055 | 2.06 | SIRIUS | GZVIFFZHYPUWML-UHFFFAOYSA-N | N-[2-(2-hydroxyethoxy)ethyl]carbamoyl fluoride |
| BRA006 | 155.082 | 2.22 | ChemWalker | OWOHLURDBZHNGG-UHFFFAOYSA-N | 2,3,6,7,8,8a-hexahydropyrrolo[1,2-a]pyrazine-1,4-dione |
| BRA006 | 111.056 | 2.42 | ChemWalker | RLOQBKJCOAXOLR-UHFFFAOYSA-N | 1H-pyrrole-2-carboxamide |
| BRA006 | 160.076 | 2.56 | ChemWalker | WHOOUMGHGSPMGR-UHFFFAOYSA-N | 2-(1H-indol-3-yl)acetaldehyde |
| BRA006 | 112.040 | 2.59 | ChemWalker | TVFIYRKPCACCNL-UHFFFAOYSA-N | furan-2-carboxamide |
| BRA006 | 164.093 | 2.63 | ChemWalker | BVIAOQMSVZHOJM-UHFFFAOYSA-N | N,N-dimethyl-7H-purin-6-amine |
| BRA006 | 111.045 | 2.95 | ChemWalker | UBTPQJCPHOHKMF-UHFFFAOYSA-N | 5-ethylidenefuran-2-one |

| | | | | | |
|---|---|---|---|---|---|
| BRA006 | 138.055 | 3.02 | GNPS2 | LOIYMIARKYCTBW-UHFFFAOYSA-N | 3-(1H-imidazol-5-yl)prop-2-enoic acid |
| BRA006 | 157.097 | 3.08 | SIRIUS | ZHDUPSJIKMSXIY-UHFFFAOYSA-N | N-methyl-2-(2-oxopyrrolidin-1-yl)acetamide |
| BRA006 | 152.071 | 3.46 | GNPS2 | CCVYRRGZDBSHFU-UHFFFAOYSA-N | 2-(2-hydroxyphenyl)acetic acid |
| BRA006 | 100.076 | 3.98 | SIRIUS | ZAEGKNQQHQDJKF-UHFFFAOYSA-O | pentanamide |
| BRA006 | 112.022 | 4.83 | SIRIUS | NIPDFDJOGCAWGH-UHFFFAOYSA-N | methoxy(methyl)borinic acid |
| BRA006 | 130.032 | 4.86 | SIRIUS | YNSUBIQKQNBKJT-UHFFFAOYSA-N | 2-(1,3-thiazol-2-yl)ethanol |
| BRA006 | 168.066 | 4.99 | GNPS2 | WKOLLVMJNQIZCI-UHFFFAOYSA-N | 4-hydroxy-3-methoxybenzoic acid |
| BRA006 | 152.071 | 6.40 | GNPS2 | SULYEHHGGXARJS-UHFFFAOYSA-N | 1-(2,4-dihydroxyphenyl)ethanone |
| BRA006 | 122.060 | 6.65 | GNPS2 | KXDAEFPNCMNJSK-UHFFFAOYSA-N | benzamide |

| | | | | | |
|---|---|---|---|---|---|
| BRA006 | 171.113 | 7.47 | ChemWalker | GINVVVBYRCSQIJ-UHFFFAOYSA-N | 2-(2-oxoazepan-1-yl)acetamide |
| BRA006 | 171.113 | 7.51 | ChemWalker | GINVVVBYRCSQIJ-UHFFFAOYSA-N | 2-(2-oxoazepan-1-yl)acetamide |
| BRA006 | 158.059 | 8.36 | ChemWalker | VUAOIXANWIFYCU-UHFFFAOYSA-N | quinoline-6-carbaldehyde |
| BRA006 | 121.065 | 9.78 | ChemWalker | BTFQKIATRPGRBS-UHFFFAOYSA-N | 2-methylbenzaldehyde |
| BRA006 | 103.054 | 9.80 | ChemWalker | UEXCJVNBTNXOEH-UHFFFAOYSA-N | ethynylbenzene |
| BRA006 | 180.102 | 9.80 | GNPS2 | ATDWJOOPFDQZNK-UHFFFAOYSA-N | N-[2-(4-hydroxyphenyl)ethyl]acetamide |
| BRA006 | 138.092 | 9.81 | ChemWalker | CDQPLIAKRDYOCB-UHFFFAOYSA-N | 4-(1-aminoethyl)phenol |
| BRA006 | 161.072 | 10.02 | GNPS2 | LSGKMZLPZFPAIN-UHFFFAOYSA-N | 1H-indole-3-carboxamide |
| BRA006 | 116.107 | 11.81 | ChemWalker | ZOFRRNUENOHELM-UHFFFAOYSA-N | 2-amino-4-methylpentanal |

| | | | | | |
|---|---|---|---|---|---|
| BRA006 | 185.129 | 11.98 | GNPS2 | JDRIJDPCYNFZIT-ACZMJKKPSA-N | (3S,6S)-3-[(2S)-butan-2-yl]-6-methylpiperazine-2,5-dione |
| BRA006 | 105.070 | 16.60 | ChemWalker | PPBRXRYQALVLMV-UHFFFAOYSA-N | styrene |
| BRA006 | 130.123 | 19.87 | ChemWalker | AEDIXYWIVPYNBI-UHFFFAOYSA-N | heptanamide |
| BRA006 | 164.106 | 19.93 | ChemWalker | LMDWIEAYQGRUHB-UHFFFAOYSA-N | 2-amino-3-phenylbutanal |
| BRA006 | 162.091 | 19.93 | ChemWalker | GLJWZMYXVRYLPB-UHFFFAOYSA-N | 3-(2-methylphenyl)prop-2-enamide |
| BRA006 | 164.107 | 20.33 | ChemWalker | NVKXBDITZDURNJ-UHFFFAOYSA-N | N-[(4-propan-2-ylphenyl)methylidene]hydroxylamine |
| BRA006 | 177.091 | 22.03 | SIRIUS | FDBYIFZQKNMIMY-UHFFFAOYSA-N | 8-fluoro-8-oxooctanoic acid |
| BRA006 | 595.381 | 30.83 | ChemWalker | IZZIZHGPMWXMBD-UHFFFAOYSA-N | 2-[[9,14-dihydroxy-14-(methoxymethyl)-3,10-dimethyl-6-propan-2-yl-8-tricyclo[9.3.0.03,7]tetradeca-1,6-dienyl]oxy]-6-(methoxymethyl)-5-(3-methylbut-1-enoxy)oxane-3,4-diol |

| | | | | | |
|---|---|---|---|---|---|
| BRA006 | 411.309 | 32.20 | ChemWalker | OELDKSOVLPNCTH-UHFFFAOYSA-N | 3-henicosa-12,16-dienoyloxy-2-hydroxypropanoic acid |
| BRA010 | 110.060 | 0.65 | ChemWalker | SOHMZGMHXUQHGE-UHFFFAOYSA-N | 5-methyl-1H-pyridin-2-one |
| BRA010 | 108.044 | 0.79 | ChemWalker | CSDSSGBPEUDDEE-UHFFFAOYSA-N | pyridine-2-carbaldehyde |
| BRA010 | 140.082 | 0.80 | ChemWalker | VYOIELONWKIZJS-UHFFFAOYSA-N | 2-amino-3-(1H-imidazol-5-yl)propanal |
| BRA010 | 199.064 | 0.82 | SIRIUS | ZYCRVTMORAHQAH-RITPCOANSA-N | [(2S,3R)-3-chloro-2-(hydroxymethyl)pyrrolidin-1-yl]-methylborinic acid |
| BRA010 | 126.055 | 0,826 | ChemWalker | VJBRLHBYLMMWER-UHFFFAOYSA-N | N-[(5-methylfuran-2-yl)methylidene]hydroxylamine |
| BRA010 | 167.093 | 0.83 | ChemWalker | PKHMCXYXGDYUJF-UHFFFAOYSA-N | 4,5,6,7-tetrahydro-3H-imidazo[4,5-c]pyridine-6-carboxamide |
| BRA010 | 185.042 | 0.89 | SIRIUS | OHCFEDLDHLJTKN-UHFFFAOYSA-N | (4-sulfinamoylphenyl)boronic acid |
| BRA010 | 113.035 | 1.10 | SIRIUS | ISAKRJDGNUQOIC-UHFFFAOYSA-N | 1H-pyrimidine-2,4-dione |

| | | | | | |
|---|---|---|---|---|---|
| BRA010 | 140.071 | 2.23 | SIRIUS | UWTVKNXZTGSDDT-UHFFFAOYSA-N | 4,6-dioxoheptanenitrile |
| BRA010 | 155.081 | 2.26 | SIRIUS | JZCWIPSHOQIMCT-UHFFFAOYSA-N | 1-(1-hydroxypropan-2-yl)imidazole-4-carbaldehyde |
| BRA010 | 146.044 | 2.73 | SIRIUS | UNEAFRQQLNNTEP-UHFFFAOYSA-N | O-(2,2,2-trifluoroethoxymethyl)hydroxylamine |
| BRA010 | 111.044 | 2.98 | SIRIUS | QIGBRXMKCJKVMJ-UHFFFAOYSA-N | benzene-1,4-diol |
| BRA010 | 199.108 | 3.03 | ChemWalker | UBLWFFBGMBRBMC-UHFFFAOYSA-N | 3-(1-hydroxyethyl)-2,3,6,7,8,8a-hexahydropyrrolo[1,2-a]pyrazine-1,4-dione |
| BRA010 | 157.097 | 3.07 | SIRIUS | MPUHTIPZENEYSH-UHFFFAOYSA-N | N-methyl-2-oxo-2-pyrrolidin-1-ylacetamide |
| BRA010 | 169.097 | 3.78 | GNPS2 | LEHOTFFKMJEONL-UHFFFAOYSA-N | 7,9-dihydro-3H-purine-2,6,8-trione |
| BRA010 | 130.032 | 4.83 | SIRIUS | ITFMBQKRDMBOON-ORCRQEGFSA-N | |
| BRA010 | 112.022 | 4.84 | SIRIUS | SCFQKZUPNGMPBM-UHFFFAOYSA-N | N-(difluoromethoxy)formamide |

| | | | | | |
|---|---|---|---|---|---|
| BRA010 | 196.097 | 6.78 | ChemWalker | OFSAJYZMIPNPHE-UHFFFAOYSA-N | N-[2-(3,4-dihydroxyphenyl)ethyl]acetamide |
| BRA010 | 137.060 | 6.83 | ChemWalker | ZWLPBLYKEWSWPD-UHFFFAOYSA-N | 2-methylbenzoic acid |
| BRA010 | 195.113 | 6.92 | ChemWalker | ASTXJGGHDAJLLS-UHFFFAOYSA-N | 4-methyl-1-(4-methyl-5-oxopyrrolidin-2-yl)-2H-pyrrol-5-one |
| BRA010 | 171.113 | 7.23 | GNPS2 | GINVVVBYRCSQIJ-UHFFFAOYSA-N | 2-(2-oxoazepan-1-yl)acetamide |
| BRA010 | 121.065 | 7.55 | SIRIUS | KWOLFJPFCHCOCG-UHFFFAOYSA-N | 1-phenylethanone |
| BRA010 | 95.086 | 8.36 | SIRIUS | HEKRPWJODWSOQV-UHFFFAOYSA-N | 3-ethenylpenta-1,4-diene |
| BRA010 | 41.091 | 8.37 | SIRIUS | PNTAWGJKWLLAAW-PHDIDXHHSA-N | (1R,6R)-bicyclo[4.1.0]heptane-7-carboxylic acid |
| BRA010 | 138.091 | 9.87 | GNPS2 | DZGWFCGJZKJUFP-UHFFFAOYSA-N | 4-(2-aminoethyl)phenol |
| BRA010 | 103.054 | 9.88 | ChemWalker | UEXCJVNBTNXOEH-UHFFFAOYSA-N | ethynylbenzene |

| | | | | | |
|---|---|---|---|---|---|
| BRA010 | 121.065 | 10.17 | ChemWalker | BTFQKIATRPGRBS-UHFFFAOYSA-N | 2-methylbenzaldehyde |
| BRA010 | 197.128 | 10.36 | GNPS2 | LGBLJRRKJGMNEN-UHFFFAOYSA-N | 1-propan-2-yl-1,2,6,7,8,8a-hexahydroimidazo[1,5-a]pyridine-3,5-dione |
| BRA010 | 178.086 | 10.54 | ChemWalker | OXFGRWIKQDSSLY-UHFFFAOYSA-N | 1,2,3,4-tetrahydroisoquinoline-1-carboxylic acid |
| BRA010 | 147.092 | 10.87 | ChemWalker | ZXIXBPGNGUZOBB-UHFFFAOYSA-N | 2-amino-2-phenylpropanenitrile |
| BRA010 | 185.128 | 12.57 | ChemWalker | COAIJFYBVHDGBZ-UHFFFAOYSA-N | 6-pentyl-1,3-diazinane-2,4-dione |
| BRA010 | 185.128 | 12.93 | GNPS2 | DBJPZCJQDRPOME-BQBZGAKWSA-N | (3S,6S)-3-methyl-6-(2-methylpropyl)piperazine-2,5-dione |
| BRA010 | 164.071 | 13.27 | ChemWalker | OWMJAQBUFVTERI-UHFFFAOYSA-N | N-(2-formylphenyl)acetamide |
| BRA010 | 164.071 | 13.83 | ChemWalker | SMIJNYOAPGHFBC-UHFFFAOYSA-N | 3-(hydroxyamino)-2-phenylprop-2-enal |
| BRA010 | 194.118 | 13.92 | ChemWalker | VIFWHZYHQSVJGD-UHFFFAOYSA-N | N-[2-(4-methoxyphenyl)ethyl]acetamide |

| | | | | | |
|---|---|---|---|---|---|
| BRA010 | 176.071 | 14.93 | GNPS2 | ZOAMBXDOGPRZLP-UHFFFAOYSA-N | 2-(1H-indol-3-yl)acetamide |
| BRA010 | 105.070 | 16.67 | ChemWalker | PPBRXRYQALVLMV-UHFFFAOYSA-N | styrene |
| BRA010 | 122.097 | 18.76 | SIRIUS | TUZXLRNKEQITEA-UHFFFAOYSA-N | N-but-3-ynylbut-2-yn-1-amine |
| BRA010 | 168.066 | 20.63 | ChemWalker | DQLYTFPAEVJTFM-UHFFFAOYSA-N | 2-amino-2-(3-hydroxyphenyl)acetic acid |
| BRA010 | 197.118 | 24.84 | ChemWalker | PWZHEFFAUWAEMU-UHFFFAOYSA-N | 3-but-2-enyl-4-(2-hydroxypropyl)-2H-furan-5-one |
| BRA010 | 109.065 | 26.84 | SIRIUS | IBZHQYDVHLIBLL-UHFFFAOYSA-N | 2-fluorobutane-1,4-diol |
| BRA010 | 431.277 | 29.12 | ChemWalker | NEZOHSNNESSNOS-UHFFFAOYSA-N | 5-(4-butan-2-yl-1,5-dihydroxy-5,6,9-trimethyl-2,4,5a,7a,8,11,11a,11b-octahydro-1H-benzo[i][3]benzoxepin-2-yl)penta-2,4-dienoic acid |
| BRA010 | 595.382 | 30.84 | ChemWalker | IZZIZHGPMWXMBD-UHFFFAOYSA-N | 2-[[9,14-dihydroxy-14-(methoxymethyl)-3,10-dimethyl-6-propan-2-yl-8-tricyclo[9.3.0.03,7]tetradeca-1,6-dienyl]oxy]-6-(methoxymethyl)-5-(3-methylbut-1-enoxy)oxane-3,4-diol |

| | | | | | |
|---|---|---|---|---|---|
| BRA010 | 639.408 | 30.87 | ChemWalker | YMIXNQVKVPYZHL-UHFFFAOYSA-N | 3-hexyl-4,6,8,10,12,14,16,27-octahydroxy-17,28-dimethyl-1-oxacyclooctacosa-17,19,21,23,25-pentaen-2-one |
| BRA010 | 439.204 | 31.89 | SIRIUS | ODGGVBKRXWOVOK-UHFFFAOYSA-N | N-[2-[2-[2-(prop-2-ynoylamino)ethoxy]ethoxy]ethyl]-11-sulfanylundecanamide |
| BRA010 | 369.239 | 31.90 | SIRIUS | JFAFCZZLYDTYHA-UHFFFAOYSA-N | [2-[[5-[(2-boronophenyl)methylamino]pentylamino]methyl]phenyl]boronic acid |
| BRA177 | 96.045 | 0.82 | ChemWalker | GCNTZFIIOFTKIY-UHFFFAOYSA-N | 1H-pyridin-4-one |
| BRA177 | 126.055 | 0.82 | ChemWalker | VJBRLHBYLMMWER-UHFFFAOYSA-N | N-[(5-methylfuran-2-yl)methylidene]hydroxylamine |
| BRA177 | 167.093 | 0.83 | SIRIUS | PVWNFAGYFUUDRC-UHFFFAOYSA-N | N-[(4-amino-2-methylpyrimidin-5-yl)methyl]formamide |
| BRA177 | 136.062 | 0.83 | GNPS2 | GFFGJBXGBJISGV-UHFFFAOYSA-N | 7H-purin-6-amine |
| BRA177 | 140.082 | 0.84 | ChemWalker | VYOIELONWKIZJS-UHFFFAOYSA-N | 2-amino-3-(1H-imidazol-5-yl)propanal |

| | | | | | |
|---|---|---|---|---|---|
| BRA177 | 108.045 | 0.84 | ChemWalker | CSDSSGBPEUDDEE-UHFFFAOYSA-N | pyridine-2-carbaldehyde |
| BRA177 | 110.060 | 0.87 | ChemWalker | OUKQTRFCDKSEPL-UHFFFAOYSA-N | 1-methylpyrrole-2-carbaldehyde |
| BRA177 | 129.066 | 1.05 | ChemWalker | LPQUIFIUJKZJRT-UHFFFAOYSA-N | 1-methyl-1,3-diazinane-2,4-dione |
| BRA177 | 113.035 | 1.07 | ChemWalker | JBTGHKUTYAMZEZ-UHFFFAOYSA-N | but-2-ynediamide |
| BRA177 | 123.055 | 1.15 | GNPS2 | DFPAKSUCGFBDDF-UHFFFAOYSA-N | pyridine-3-carboxamide |
| BRA177 | 137.046 | 1.29 | GNPS2 | FDGQSTZJBFJUBT-UHFFFAOYSA-N | 3,7-dihydropurin-6-one |
| BRA177 | 153.041 | 1.48 | GNPS2 | LRFVTYWOQMYALW-UHFFFAOYSA-N | 3,7-dihydropurine-2,6-dione |
| BRA177 | 113.035 | 1.84 | SIRIUS | ISAKRJDGNUQOIC-UHFFFAOYSA-N | 1H-pyrimidine-2,4-dione |
| BRA177 | 127.050 | 1.93 | GNPS2 | RWQNBRDOKXIBIV-UHFFFAOYSA-N | 5-methyl-1H-pyrimidine-2,4-dione |

| | | | | | |
|---|---|---|---|---|---|
| BRA177 | 136.062 | 1.94 | ChemWalker | GFFGJBXGBJISGV-UHFFFAOYSA-N | 7H-purin-6-amine |
| BRA177 | 174.055 | 2.04 | ChemWalker | PCPFJNLCUUEVDQ-UHFFFAOYSA-N | 4-oxo-1H-quinoline-2-carbaldehyde |
| BRA177 | 157.097 | 2.22 | GNPS2 | GFFGJBXGBJISGV-UHFFFAOYSA-N | 7H-purin-6-amine |
| BRA177 | 155.081 | 2.23 | ChemWalker | GRDXZRWCQWDLPG-UHFFFAOYSA-N | 1,3,6-trimethylpyrimidine-2,4-dione |
| BRA177 | 111.044 | 2.41 | ChemWalker | UBTPQJCPHOHKMF-UHFFFAOYSA-N | 5-ethylidenefuran-2-one |
| BRA177 | 164.093 | 2.54 | ChemWalker | BVIAOQMSVZHOJM-UHFFFAOYSA-N | N,N-dimethyl-7H-purin-6-amine |
| BRA177 | 160.076 | 2.60 | SIRIUS | WHOOUMGHGSPMGR-UHFFFAOYSA-N | 2-(1H-indol-3-yl)acetaldehyde |
| BRA177 | 199.108 | 2.92 | GNPS2 | UBLWFFBGMBRBMC-UHFFFAOYSA-N | 3-(1-hydroxyethyl)-2,3,6,7,8,8a-hexahydropyrrolo[1,2-a]pyrazine-1,4-dione |
| BRA177 | 157.097 | 3.02 | ChemWalker | QOSQDHSPKMBGMH-UHFFFAOYSA-N | 2-(2-oxopiperidin-1-yl)acetamide |

| | | | | | |
|---|---|---|---|---|---|
| BRA177 | 169.097 | 3.73 | GNPS2 | LEHOTFFKMJEONL-UHFFFAOYSA-N | 7,9-dihydro-3H-purine-2,6,8-trione |
| BRA177 | 127.050 | 3.92 | GNPS2 | HDFGOPSGAURCEO-UHFFFAOYSA-N | 1-ethylpyrrole-2,5-dione |
| BRA177 | 125.071 | 3.95 | ChemWalker | GCQHUBANENYTLB-UHFFFAOYSA-N | 2-(1-methylimidazol-4-yl)acetaldehyde |
| BRA177 | 130.032 | 4.77 | ChemWalker | MWSQHVUUIHWHBM-UHFFFAOYSA-N | 2-(1,3-thiazol-5-yl)ethanol |
| BRA177 | 112.040 | 4.84 | ChemWalker | ZLPORNPZJNRGCO-UHFFFAOYSA-N | 3-methylpyrrole-2,5-dione |
| BRA177 | 195.113 | 6.82 | ChemWalker | KIPFMNAHRGYDMB-UHFFFAOYSA-N | 2-(aminomethyl)-5-methoxy-N-methylbenzamide |
| BRA177 | 171.113 | 7.18 | ChemWalker | GINVVVBYRCSQIJ-UHFFFAOYSA-N | 2-(2-oxoazepan-1-yl)acetamide |
| BRA177 | 146.060 | 7.47 | ChemWalker | OLNJUISKUQQNIM-UHFFFAOYSA-N | 1H-indole-3-carbaldehyde |
| BRA177 | 171.112 | 7.55 | SIRIUS | HPHUVLMMVZITSG-UHFFFAOYSA-N | 2-(2-oxopyrrolidin-1-yl)butanamide |

| | | | | | |
|---|---|---|---|---|---|
| BRA177 | 103.054 | 9.73 | GNPS2 | RGHHSNMVTDWUBI-UHFFFAOYSA-N | 4-hydroxybenzaldehyde |
| BRA177 | 121.065 | 9.75 | ChemWalker | FUGYGGDSWSUORM-UHFFFAOYSA-N | 4-ethenylphenol |
| BRA177 | 138.091 | 9.75 | GNPS2 | DZGWFCGJZKJUFP-UHFFFAOYSA-N | 4-(2-aminoethyl)phenol |
| BRA177 | 197.128 | 10.25 | GNPS2 | LGBLJRRKJGMNEN-UHFFFAOYSA-N | 1-propan-2-yl-1,2,6,7,8,8a-hexahydroimidazo[1,5-a]pyridine-3,5-dione |
| BRA177 | 169.076 | 11.96 | ChemWalker | JIFVPBALUHDGEP-UHFFFAOYSA-N | 2-[(4-methylphenyl)methylidene]propanedinitrile |
| BRA177 | 185.128 | 11.98 | GNPS2 | COAIJFYBVHDGBZ-UHFFFAOYSA-N | 6-pentyl-1,3-diazinane-2,4-dione |
| BRA177 | 185.128 | 12.78 | GNPS2 | DBJPZCJQDRPOME-BQBZGAKWSA-N | (3S,6S)-3-methyl-6-(2-methylpropyl)piperazine-2,5-dione |
| BRA177 | 164.071 | 13.66 | ChemWalker | SMIJNYOAPGHFBC-UHFFFAOYSA-N | 3-(hydroxyamino)-2-phenylprop-2-enal |
| BRA177 | 176.071 | 14.91 | GNPS2 | ZOAMBXDOGPRZLP-UHFFFAOYSA-N | 2-(1H-indol-3-yl)acetamide |

| | | | | | |
|---|---|---|---|---|---|
| BRA177 | 91.054 | 17.04 | ChemWalker | AMSMVCOBCOZLEE-UHFFFAOYSA-N | bicyclo[4.1.0]hepta-1,3,5-triene |
| BRA177 | 137.060 | 17.11 | GNPS2 | RWZYAGGXGHYGMB-UHFFFAOYSA-N | 2-aminobenzoic acid |
| BRA177 | 122.097 | 18.47 | GNPS2 | WPYMKLBDIGXBTP-UHFFFAOYSA-N | benzoic acid |
| BRA177 | 105.070 | 18.71 | ChemWalker | PPBRXRYQALVLMV-UHFFFAOYSA-N | styrene |
| BRA177 | 164.107 | 18.73 | GNPS2 | MODKMHXGCGKTLE-UHFFFAOYSA-N | N-(2-phenylethyl)acetamide |
| BRA177 | 187.089 | 21.09 | ChemWalker | CXBYCIKWTYCLAR-UHFFFAOYSA-N | 1-methyl-5-phenylpyrimidin-2-one |

Supplementary Table 2. Fragmented ions belonging exclusively to the list (SPL) that presented the annotation and their respective metabolic pathways.

| Sample | Compound (IUPAC name) | Metabolic pathway (KEGG) | KEGG compound ID | KEGG pathway ID |
|---|---|---|---|---|
| BRA006 | 2-methylbenzaldehyde | Xylene degradation | cpd:C07214 | map00622 |
| BRA006 | benzamide | Aminobenzoate degradation | cpd:C09815 | map00627 |
| BRA006 | 3,7-dihydropurine-2,6-dione | Purine metabolism | cpd:C00385 | map00230 |
| BRA006 | 3-(1H-imidazol-5-yl)prop-2-enoic acid | Histidine metabolism | cpd:C00785 | map00340 |
| BRA006 | 2-(2-hydroxyphenyl)acetic acid | Phenylalanine metabolism | cpd:C05852 | map00360 |
| BRA006 | styrene | Ethylbenzene degradation | cpd:C07083 | map00642 |
| BRA006 | 4-hydroxy-3-methoxybenzoic acid | Aminobenzoate degradation | cpd:C06672 | map00627 |
| BRA006 | 2-(1H-indol-3-yl)acetaldehyde | Tryptophan metabolism | cpd:C00637 | map00380 |
| BRA006 | N-[(4-amino-2-methylpyrimidin-5-yl)methyl]formamide | Thiamine metabolism | cpd:C19872 | map00730 |
| BRA010 | 1H-pyrimidine-2,4-dione | Pyrimidine metabolism | cpd:C00106 | map00240 |
| BRA010 | (i) 4-(2-aminoethyl)phenol ; (ii) benzene-1,4-diol | Tyrosine metabolism | (i) cpd:C00483; (ii) cpd:C00530 | map00350 |
| BRA010 | styrene | Styrene degradation | cpd:C07083 | map00643 |

| | | | | |
|---|---|---|---|---|
| BRA010 | 2-(1H-indol-3-yl)acetamide | Tryptophan metabolism | cpd:C02693 | map00380 |
| BRA010 | (i) 2-methylbenzoic acid; (ii) 2-methylbenzaldehyde | Xylene degradation | (i) cpd:C07215; (ii) cpd:C07214 | map00622 |
| BRA010 | 2-amino-3-(1H-imidazol-5-yl)propanal | Histidine metabolism | cpd:C01929 | map00340 |
| BRA010 | 1-phenylethanone | Ethylbenzene degradation | cpd:C07113 | map00642 |
| BRA010 | 7,9-dihydro-3H-purine-2,6,8-trione | Purine metabolism | cpd:C00366 | map00230 |
| BRA177 | (i) 7H-purin-6-amine; (ii) 3,7-dihydropurin-6-one; (iii) 3,7-dihydropurine-2,6-dione; (iv) 7,9-dihydro-3H-purine-2,6,8-trione | Purine metabolism | (i) cpd:C00147; (ii): cpd:C00262; (iii) cpd:C00385; (iv) cpd:C00366 | map00230 |
| BRA177 | 4-ethenylphenol | Phenylpropanoid biosynthesis | cpd:C05627 | map00940 |
| BRA177 | 4-(2-aminoethyl)phenol | Tyrosine metabolism | cpd:C00483 | map00350 |
| BRA177 | styrene | Styrene degradation | cpd:C07083 | map00643 |
| BRA177 | (i) 5-methyl-1H-pyrimidine-2,4-dione; (ii) 1H-pyrimidine-2,4-dione | Pyrimidine metabolism | (i) cpd:C00178; (ii) cpd:C00106 | map00240 |
| BRA177 | N-[(4-amino-2-methylpyrimidin-5-yl)methyl]formamide | Thiamine metabolism | cpd:C19872 | map00730 |

| | | | | |
|---|---|---|---|---|
| BRA177 | (i) 2-(1H-indol-3-yl)acetaldehyde; (ii) 2-aminobenzoic acid; (iii) 2-(1H-indol-3-yl)acetamide | Tryptophan metabolism | (i) cpd:C00637; (ii) cpd:C00108; (iii) cpd:C02693 | map00380 |
| BRA177 | (i) 2-(1-methylimidazol-4-yl)acetaldehyde; (ii) 2-amino-3-(1H-imidazol-5-yl)propanal | Histidine metabolism | (i) cpd:C05827; (ii) cpd:C01929 | map00340 |
| BRA177 | 4-hydroxybenzaldehyde | Aminobenzoate degradation | cpd:C00633 | map00627 |
| BRA177 | pyridine-3-carboxamide | Nicotinate and nicotinamide metabolism | cpd:C00153 | map00760 |
| BRA177 | benzoic acid | Benzoate degradation | cpd:C00180 | map00362 |
